# CXCR3-CXCL11 signaling restricts angiogenesis and promotes pericyte recruitment

**DOI:** 10.1101/2023.09.16.557842

**Authors:** Megan E. Goeckel, Jihui Lee, Allison Levitas, Sarah Colijn, Geonyoung Mun, Zarek Burton, Bharadwaj Chintalapati, Ying Yin, Javier Abello, Amber Stratman

**Affiliations:** Department of Cell Biology and Physiology, Washington University School of Medicine St. Louis, MO, 63110; University of Nebraska Medical Center, Graduate Studies, Nebraska Medical Center, Omaha, NE 68198

**Keywords:** endothelial cells, pericyte, cardiovascular development, chemokine signaling

## Abstract

Endothelial cell (EC)-pericyte interactions are known to remodel in response to hemodynamic forces, yet there is a lack of mechanistic understanding of the signaling pathways that underlie these events. Here, we have identified a novel signaling network regulated by blood flow in ECs—the chemokine receptor, CXCR3, and one of its ligands, CXCL11—that delimits EC angiogenic potential and suppresses pericyte recruitment during development through regulation of *pdgfb* expression in ECs. *In vitro* modeling of EC-pericyte interactions demonstrates that suppression of EC-specific CXCR3 signaling leads to loss of pericyte association with EC tubes. *In vivo*, phenotypic defects are particularly noted in the cranial vasculature, where we see a loss of pericyte association with and expansion of the vasculature in zebrafish treated with the Cxcr3 inhibitor AMG487. We also demonstrate using flow modeling platforms that CXCR3-deficient ECs are more elongated, move more slowly, and have impaired EC-EC junctions compared to their control counterparts. Together these data suggest that CXCR3 signaling in ECs drives vascular stabilization events during development.

## INTRODUCTION

The blood vessel wall is composed of both endothelial cells (EC) and mural cells (MC)—such as vascular smooth muscle cells and pericytes ^1^. Throughout life, proper formation and stabilization of the vascular wall is critical for halting vessel outgrowth and preventing vessel leak/hemorrhage. This is accomplished by communication between ECs and MCs. MCs have been shown to localize to EC-EC junctions to maintain vascular barrier function, and EC-MC interactions have been shown to be critical during the formation of the vascular basement membrane ^2-6^. As the vasculature constantly experiences hemodynamic forces, coordinated cell-to-cell interactions are critical to facilitate proper vascular development ^7^. For instance, during development, MCs are known to preferentially associate with arteries ^8^; however, while this structural adaptation is critical for stabilizing blood vessels to withstand sheer and radial stress, the mechanisms required for localization of MCs specifically to arteries is still being explored. Arterial localization of MCs is dependent on the activation state of Krüppel-like transcription factor 2 (KLF2), a zinc-finger transcription factor regulated by blood flow ^9^. During early embryogenesis, KLF2 is highly expressed in venous endothelium of zebrafish and mice, functioning as a transcriptional repressor of the pro-MC recruitment molecule platelet derived growth factor subunit B (PDGFB) ^9^. Therefore, suppressing KFL2 levels in the venous endothelium leads to increased PDGFB production and aberrant MC localization ^9^. These findings show that mechanical cues to blood flow are essential for genetic and cellular behavior. Although blood flow plays an important role in the formation and remodeling of vascular networks during development, the cellular and molecular mechanisms remain elusive.

In immune cells, KLF2 activity has been shown to indirectly repress expression of proinflammatory chemokine receptors such as CXC motif chemokine receptor 3 (CXCR3)^10^. Chemokine ligands—which are small, soluble proteins—interact with the CXC family of G-protein-coupled receptors to attract leukocytes to sites of inflammation ^11-13^. While classically described for their role in the immune response, members of the CXC family are known to be expressed in endothelium ^14^. CXC motif chemokine receptor 4 (CXCR4), and its ligand CXC motif chemokine ligand 12 (CXCL12) have known roles in promoting angiogenesis ^15^ and—as described in our previous work—contributing to MC recruitment through upregulation of PDGFBB expression on arteries ^9^.

Alternatively, certain CXC family chemokines have been suggested to inhibit angiogenesis ^16^. The receptor CXCR3 and its ligand, CXC motif chemokine ligand 11 (CXCL11), have been shown to have a negative effect on EC motility *in vitro* ^17^. CXCR3 has been described as an angiostatic chemokine receptor and lacks a sequence of amino acids (Glu-Leu-Arg, “ELR”) on the N-terminus that seems to control motility in endothelial cells ^18^. Several studies highlight that chemokines and chemokine receptors affect vascular development—including angiogenesis, lymphangiogenic potential differentiation, and recruitment of MCs ^19-24^. These suggest the importance of chemokine signaling in the regulation of vascular function and structure through communication between ECs and MCs. Surprisingly, virtually nothing is known about the role of CXCR3 signaling in regulating early vascular development and vessel wall stabilization events. Therefore, in this study, we aim to characterize the role of CXCR3 in regulating blood vessel formation and pericyte recruitment during early vascular patterning events.

Here, we use 3-dimensional (3D) collagen matrix assays to demonstrate EC tube formation *in vitro* and embryonic zebrafish to model these findings *in vivo.* Upon CXCR3 activation, EC tube formation is decreased *in vitro* and zebrafish cranial vascular patterning is negatively affected *in vivo*. Upon CXCR3 inhibition, we observe increased EC tube formation *in vitro* and expansion of the zebrafish cranial vascular network. EC-specific CXCR3 inhibition also leads to a loss recruitment and differentiation of pericytes around developing EC tube networks in 3D collagen matrix assays, which is replicated *in vivo* in the zebrafish cranial vasculature. Using flow-modeling assays, we also show that CXCR3-deficient ECs are more elongated with decreased cell motility and impaired EC-EC junctions. Therefore, this work demonstrates a novel role for CXCR3 in restricting vascular outgrowth and promoting vessel stabilization via regulation of cell motility, EC junctions, and pericyte association with the vasculature.

## RESULTS

### CXCR3 is expressed in endothelial cells during cardiovascular development and is regulated by hemodynamic forces

Through mining of published single cell sequencing data from different developmental stages in zebrafish ^25^, we sought to confirm *cxcr3* expression in the developing endothelium by tracking early mesodermal lineages that can differentiate into endothelial cells—such as the heart primordium cells and the intermediate cell mass cells (ICM) (**Fig. 1A,B**). The distribution (i.e. localization of cells in the data set) of heart primordium and ICM cell lineages are shown superimposed on the temporal coded UMAP (**Fig. 1A,B**). There are three *cxcr3* isoforms in the zebrafish—*cxcr3.1*, *cxcr3.2*, and *cxcr3.3*. UMAP plots show a variety of cells expressing *cxcr3.1* and *cxcr3.3* across these embryonic stages (*cxcr3.2* is not expressed at high enough levels in this dataset to have been annotated, **Fig. 1C**). To identify the cell populations that express *cxcr3.1* and *cxcr3.3*, we assembled UMAP plots of genes that mark cardiovascular and hematopoietic cells derived from the early mesoderm—endothelial cells (*ephb2b, chd5, kdrl, fli1a, klf2a, and cxcr4a*), blood lineages (*tal1, lmo2, gata1a, fzd5, and cxcr4a*), and cardiomyocytes (*myl6, tbx5a, nkx2.5, cxcr4a*, and *sgcb*)—demonstrating that the *cxcr3.1* isoform has a gene expression profile that most similarly tracks endothelial cell lineage markers, and *cxcr3.3* is broadly dispersed (**Fig. 1C; Supp. Fig. 1**).

**Figure 1.**
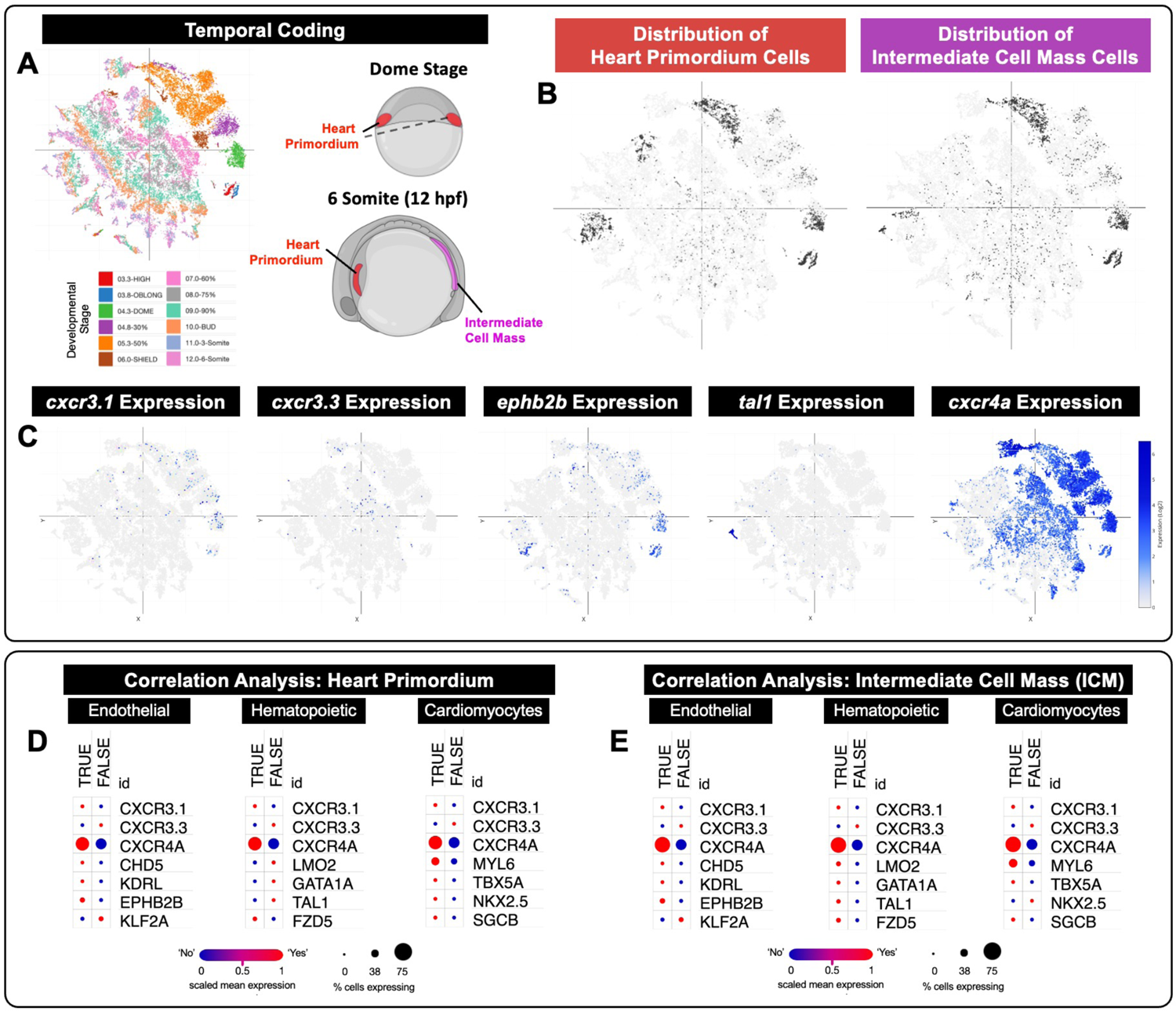
Single cell analysis of zebrafish embryos reveals expression of *cxcr3* isoforms in the heart primordium and intermediate cell mass during development. A) Using publicly available single cell sequencing data sets from Farrell, et. al. 2018 ^25^ and Sur, et. al. 2023 ^26^, we show a UMAP plot of cell populations temporally coded across developmental stages from the Dome Stage to the 6-somite stage. Cartoons of dome stage and 6 somite zebrafish embryos show where our cell lineages of interest are located anatomically at each stage. Red, heart primordium; Magenta, intermediate cell mass. **B)** Distribution (i.e. location) of heart primordium cells and intermediate cell mass cells (dark grey dots) within the temporal coded UMAP plot (light grey dots). **C)** UMAP plots of gene expression (blue dots) shows individual cells that express *cxcr3.1*, *cxcr3.3, eph2b2* (endothelial cell marker), *tal1* (hematopoietic lineage marker), and *cxcr4a* (broad endothelial, hematopoietic, and neuronal marker) across all developmental stages. Darker blue dots equate to higher gene expression, gray dots equate to no gene expression. **D,E)** Correlation analysis of markers that define the heart primordium (D) and the intermediate cell mass (E) with *cxcr3.1*, *cxcr3.3* and known endothelial, hematopoietic, and cardiomyocyte markers. Red color indicates a positive correlation, blue color indicates a negative correlation; size of the dot corresponds to the percent of cells in a given lineage that express the indicated gene. As shown, endothelial and cardiomyocyte markers correlate with the heart primordium, suggesting that these two cell types derive from this precursor population. Endothelial, hematopoietic, and some cardiomyocyte markers correlate with the intermediate cell mass, suggesting all three lineages can arise from these precursors.

We next used this data to generate correlation analysis between individual gene expression profiles and cellular lineages ^25,26^. We analyzed the co-expression of *cxcr3.1* with known endothelial, hematopoietic, and cardiomyocyte cell markers within the heart primordium or the ICM cell populations (**Fig. 1D,E**). These analyses show that the heart primordium generates cell populations that express markers of and become endothelial cells (*cxcr4a, chd5, kdrl, ephb2b*) and cardiomyocytes (*cxcr4a, myl6, tbx5a, nkx2.5, sgcb*). Within this subset of heart primordium cells, *cxcr3.1* expression positively correlates with expression of the genes marking both endothelial cells and cardiomyocytes (i.e. red dots under ‘True’; **Fig. 1D**). The same analysis was done using ICM cells. The ICM generates cell populations that express markers of and become endothelial cells (*cxcr4a, chd5, kdrl, ephb2b*), hematopoietic cells (*cxcr4a, lmo2, gata1a, tal1, fzd5*), and possibly some cardiomyocytes (*cxcr4a, myl6, tbx5a, sgcb*, but not *nkx2.5*). Within this sub-population of ICM cells, *cxcr3.1* expression positively correlates with markers of all three lineages, endothelial, hematopoietic, and cardiomyocytes (**Fig. 1E**). Taken together, this suggests that there are early mesoderm cell populations that express *cxcr3.1* in the zebrafish, and these cells can give rise to cardiovascular or hematopoietic lineages. However, a large percentage of *cxcr3.1* positive cells co-express markers of endothelial cells at these early developmental stages.

We carried out similar analysis by mining published E8.25 mouse single cell sequencing data ^27^, comparing UMAP plots of *Cxcr3* expression to markers of endothelial cells (*Kdr*, *Ephb2*, *Fli1*, *Cxcr4*, and *Klf2*), hematopoietic cells (*Gata1*, *Lmo2*, *Tal1*, *Cxcr4*, and *Fzd5*), and cardiomyocytes (*Myl7*, *Tbx5*, *Nkx2.5*, *Cxcr4*, and *Sgca*) (**Fig. 2; Supp. Fig. 2**). Cell type clustering from this dataset is published and identifies endothelium, embryonic blood, and cardiac populations (**Fig. 2A**) ^27^. When analyzing UMAP analyses of endothelial genes versus cell clusters, *Cxcr3* is shown to be almost exclusively expressed in the endothelium at this stage of murine development (**Fig. 2B; Supp. Fig. 2**), with minimal *Cxcr3* expression that overlaps with embryonic blood/hematopoietic markers or cardiac/cardiomyocytes markers (**Supp. Fig. 2**).

**Figure 2.**
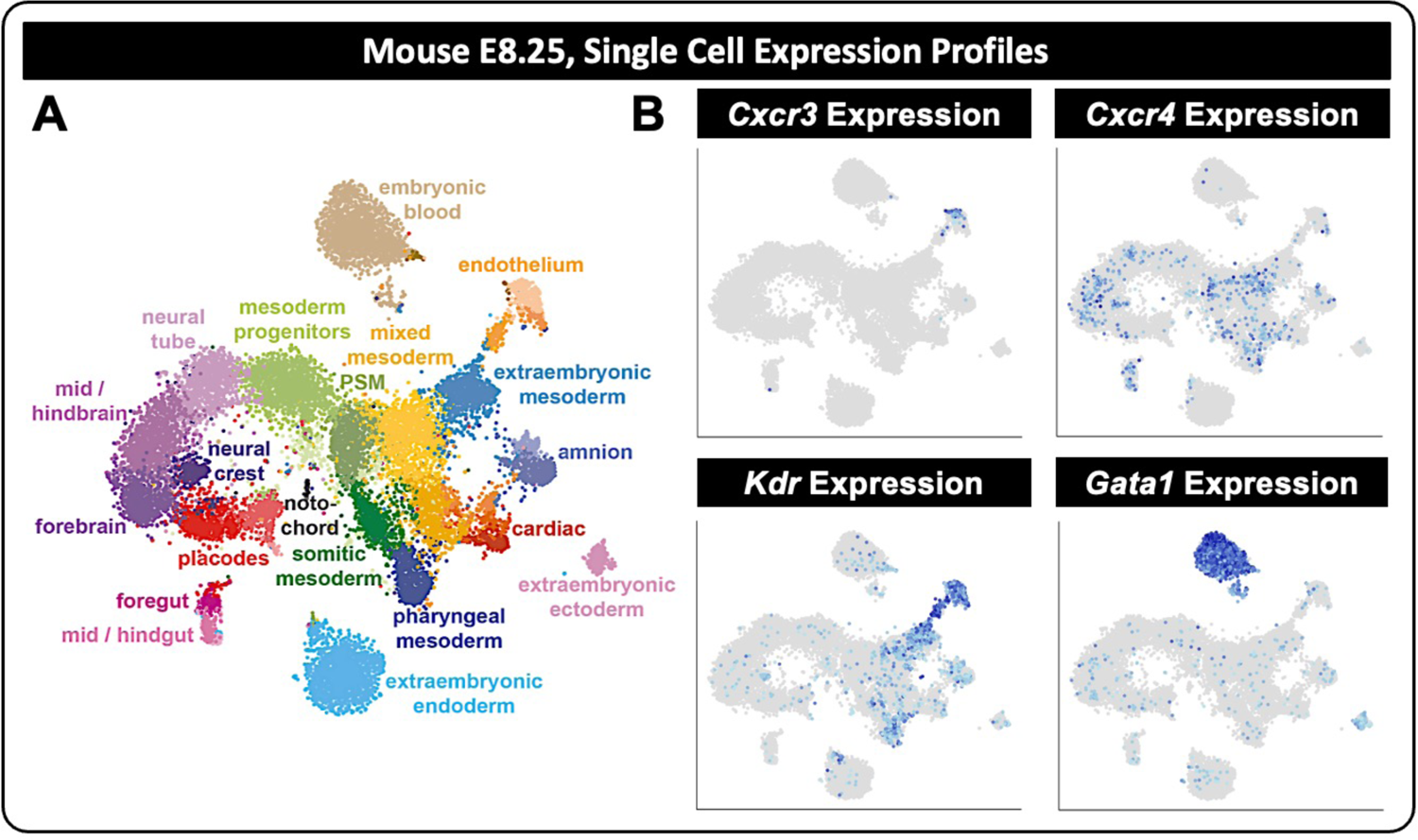
Single cell analysis of E8.25 mouse embryos reveals expression of *CXCR3* in endothelial cells during development. **A)** Using publicly available single cell sequencing data sets from Ibarra-Soria, et. al. 2018 ^27^, we show a UMAP plot clustering cell populations from the mouse E8.25 embryo. **B)** UMAP plots of gene expression (blue dots) shows individual cells that express *Cxcr3, Kdr* (endothelial cell marker), *Gata1* (hematopoietic lineage marker), and *Cxcr4* (broad endothelial, hematopoietic, and neuronal marker) at E8.25. Darker blue dots equate to high gene expression, gray dots equate to no gene expression. As shown, at E8.25 in the mouse, *Cxcr3* is almost exclusively restricted to the endothelium.

To confirm *cxcr3* isoform expression in zebrafish endothelial cells and determine if expression of *cxcr3* isoforms are responsive to changes in shear stress, we treated zebrafish embryos with BDM (2,3-Butanedione 2-monoxime, a myosin ATPase inhibitor) to inhibit cardiac muscle contraction ^28^. We then performed EC-specific mRNA isolation via translational ribosomal affinity purification (TRAP) followed by mRNA sequencing or qPCR (**Supp. Fig. 3**, **Fig. 3A-D**); or we fixed samples for *in situ* hybridization. *klf2a* transcript, which has been reported to decrease when blood flow is suppressed, is shown as an experimental control, confirming that we were able to sufficiently suppress blood flow in these samples (**Fig. 3A**). We next looked at expression of *cxcl11* and *cxcr3* isoforms in these samples. *cxcl11*—which has five tandem transcripts in the zebrafish—is the primary ligand for *cxcr3* present in ECs at these stages ^29^. mRNA sequencing of *cxcl11.1, cxcl11.5, cxcl11.6, cxcl11.7, and cxcl11.8* transcripts, and confirmatory *in situ* hybridization of *cxcl11.1*, shows that the *cxcl11* class of ligands is upregulated in ECs by suppression of blood flow (**Fig. 3B,E**). Additionally, we see significant upregulation of the *cxcr3.1* transcript in ECs following suppression of blood flow (**Fig. 3C,E**). We used *in situ* hybridization to confirm localization of the upregulated *cxcr3.1* transcripts. Under normal blood flow conditions (**Fig. 3E**, Flow), *cxcr3.1* transcript is essentially not expressed by 48 hpf; however, when blood flow is suppressed (**Fig. 3E**, No Flow) *cxcr3.1* transcript is maintained at a high level in the vasculature. In the zebrafish, the *cxcr3.2* isoform is reported to be a signaling ‘trap’, working in opposition to *cxcr3.1*. In support of the early-stage zebrafish single cell sequencing analysis, we do not see *cxcr3.2* expression in the endothelium at 48 hpf by *in situ* hybridization, nor significant regulation of *cxcr3.2* by blood flow in our EC-specific TRAP mRNA isolation and sequencing analysis (**Fig. 3C,F**). Finally, while *cxcr3.3* was only modestly expressed in the zebrafish single cell dataset, we show by EC-specific TRAP mRNA isolation and qPCR analysis that it can be enriched in the endothelium following suppression of blood flow (**Fig. 3D**). Taken together, these data strongly support the finding that endothelial cells express *cxcr3.1, cxcr3.3,* and *cxcl11* isoforms during development—without significant levels of *cxcr3.2* being present—and that *cxcr3.1, cxcr3.3,* and *cxcl11* transcript levels can be regulated by changes in hemodynamics.

**Figure 3.**
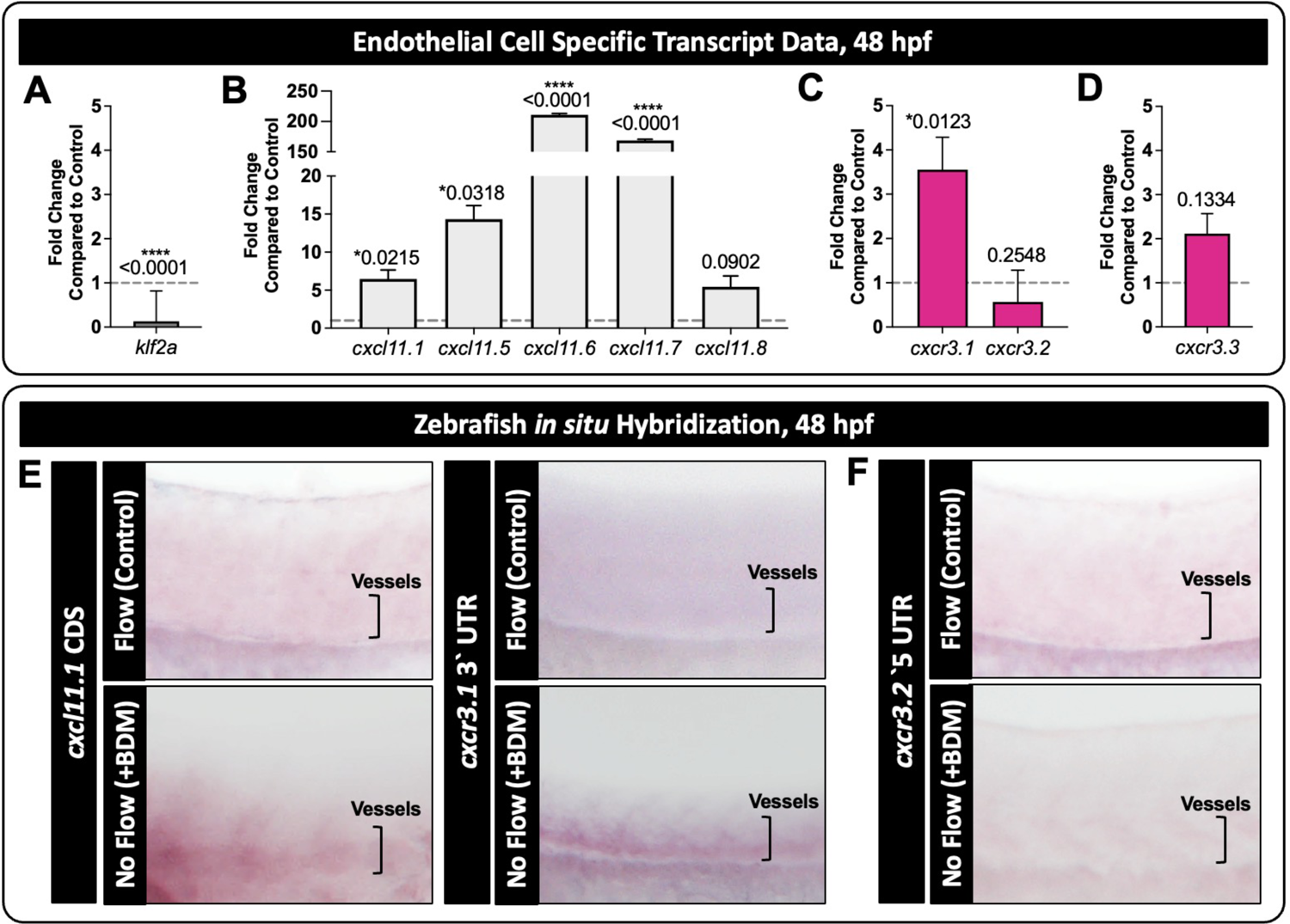
*cxcr3.1, cxcr3.3,* and *cxcl11* isoforms are expressed in endothelial cells and their transcripts are responsive to blood flow. **A-C)** mRNA isolated via the TRAP protocol (Supp. Fig. 3) and then mRNA-sequenced was used to assess EC-specific transcript levels of *klf2a* (A), *cxcl11.1*, *cxcl11.5*, *cxcl11.6*, *cxcl11.7,* and *cxcl11.8* (B), *cxcr3.1,* and *cxcr3.2* (C) with (Control) and without (+BDM) blood flow. The data is presented as relative fold change of +BDM compared to the control (grey dotted line). **D)** *cxcr3.3* was not represented in the mRNA sequencing data set and therefore was assessed by targeted qPCR. Data is reported in the same manner as A-C. **E,F)** Representative images of *in situ* hybridization from 48 hpf zebrafish embryos. Flow was either allowed to start normally (Flow) or inhibited by the addition of BDM at 22 hpf (No Flow). Location of the blood vessels are shown in the brackets, purple color marks RNA transcripts. Both *cxcl11.1* and *cxcr3.1* transcripts are increased in response to impaired blood flow (E), while *cxcr3.2* does not change (F). Data in A-C are two RNA-Seq replicates of 3,000 embryos each and are presented as bar graphs that display the mean±SEM. The mean, SEM, and p-values were reported from the RNA-Seq dataset. Data in D are presented as bar graphs that display the mean±SD from n=3 independent mRNA isolations and statistics were calculated using a one-sample t test.

### CXCR3 is required *in vivo* to limit cranial vascular angiogenesis in the zebrafish during development

To address the gain- and loss-of-function roles of CXCR3 on vascular development *in vivo*, we used a series of pharmacologic reagents to modulate CXCR3 activity—the agonist (VUF11222) and the antagonist (AMG487). We treated *Tg(fli1a:eGFP)* embryos with AMG487 and VUF11222 at 32 hpf, when the cranial vasculature is beginning to expand, and analyzed phenotypes at 54 hpf. Confocal imaging reveals that suppression of CXCR3 activity (AMG487) promotes expansion of the vascular network, while activation of CXCR3 activity (VUF11222) inhibits expansion of the vascular network (**Fig. 4A-C**). Total vascular area, as quantified by automated GFP-masking, reveals an increase in vascular area in the AMG487 treatment condition and a decrease in vascular area in the VUF11222 treatment condition (**Fig. 4D**). Average number of branch points in the vessels trend in the same directions as vascular area, though do not reach statistical significance (**Fig. 4E**). We note no changes in the length of individual vessel segments compared to the vertical radius of the cranial space (**Fig. 4F**). Whole-mount immunostaining for cell motility regulator, phosphorylated extracellular signal-regulated kinase 1/2 (p-ERK1/2), reveals an increase in expression of p-ERK1/2 when CXCR3 activity is inhibited with AMG487 (magenta; **Fig. 4G,H**). We note minimal effects of CXCR3 activation with VUF11222 on p-ERK1/2.

**Figure 4.**
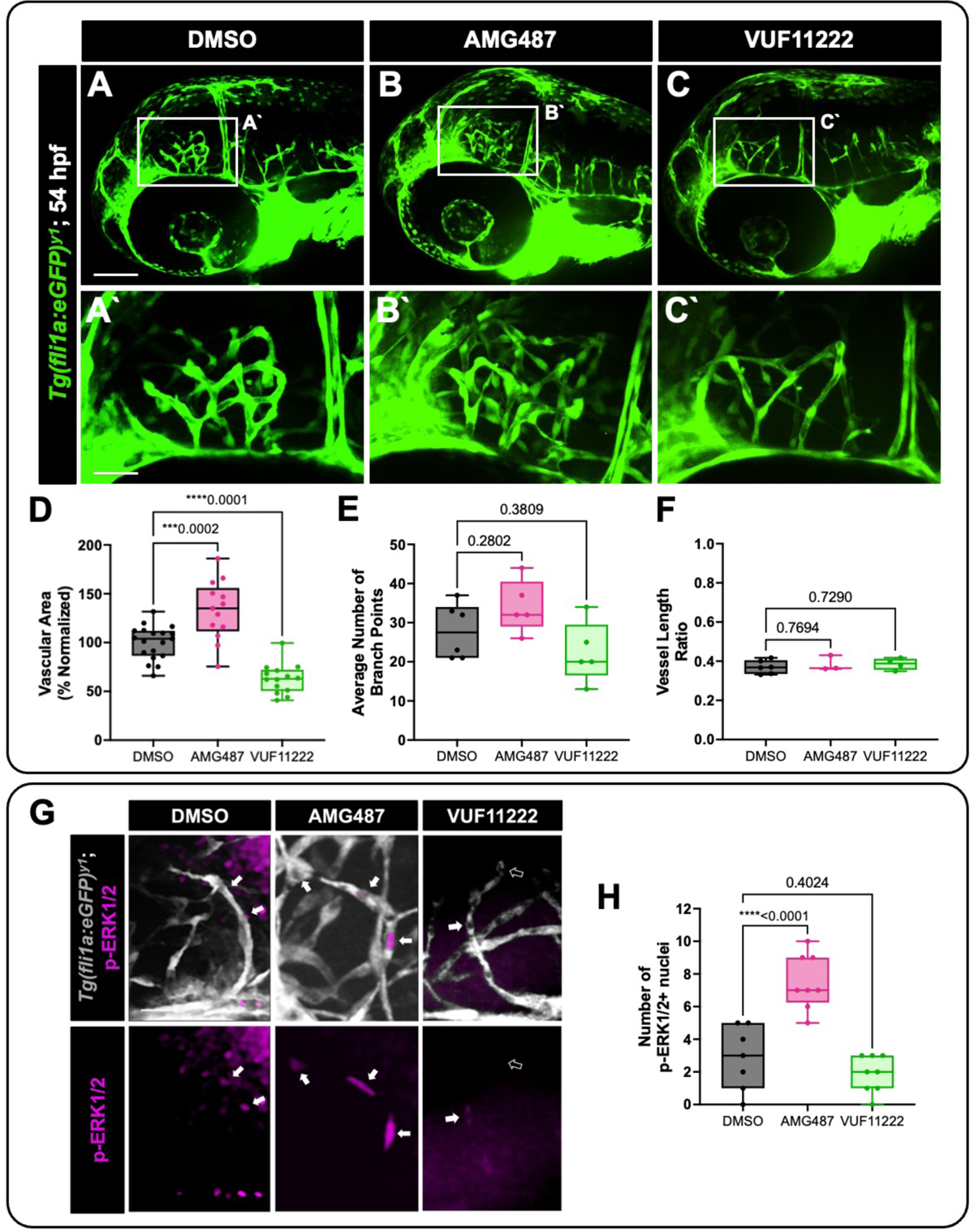
CXCR3 contributes to cranial angiogenesis in the developing zebrafish. **A-C)** Representative images of *Tg(fli1a:GFP)* zebrafish treated with DMSO vehicle control (A), the CXCR3 antagonist (AMG487, B), or the CXCR3 agonist (VUF11222, C). Pharmacological reagents were added to the zebrafish water at 32 hpf and embryos were imaged at 54 hpf to view changes in vascularization. Panels A’-C’ show magnified insets. Scale bars represent 500µm (A-C) and 50µm (A’-C’). **D)** Quantification of vascular area, normalized to the DMSO control. **E)** Average number of EC vessel branch points in the frontal lobe of the zebrafish brain (the region shown in the images). **F)** Average vessel length. To account for any changes in mounting angle of the samples, a vessel length ratio was calculated by measuring the length of each vessel and then dividing by the distance from the top of the eye to the top of the skull for each fish. **G)** Representative images of whole mount immunofluorescent staining of p-ERK1/2 in DMSO control, AMG487, and VUF11222 treated *Tg(fli1a:GFP)* zebrafish (vessels in grey). White arrows show positive p-ERK1/2 immunostaining (magenta), while the open arrows show p-ERK1/2-negative nuclei. Scale bar represents 25µm. **H)** Quantification of the number of p-ERK1/2-positive nuclei within the vasculature of the frontal lobe of the zebrafish (region shown in A’-C’). For panels D, E, F, and H, each dot represents an individual zebrafish. One outlier was identified and removed from the VUF11222 group in panel D as per the ROUT method (Q=1%). Statistics for these panels were calculated using one-way ANOVA with Dunnett’s multiple comparisons test. Omnibus ANOVA *P*-values (prior to the *post hoc* tests) are <0.0001 (D), 0.0649 (E), 0.729 (F), and <0.0001 (H). Data are presented as box plots that display the median value with 1^st^ and 3^rd^ quartiles and min/max bars.

### CXCR3 regulates endothelial cell motility and junctional stabilization *in vitro*

To assess the functional role of CXCR3 in EC biology at a mechanistic level, we utilized two *in vitro* platforms to test 1) EC cellular responses to flow and 2) the ability of ECs to form tubes and recruit pericytes, which model vessel formation and stabilization events. As we show CXCR3 is a blood flow responsive gene, we wanted to determine if ECs utilize the gene for interpreting flow forces and/or flow-mediated signaling. To determine this, we used our recently developed blood flow modeling platform, where we can seed ECs as a monolayer and then provide various flow types across the surface of the cells to mimic the shear stresses experienced in vivo ^30^. We compared siRNA-treated control ECs (siControl) to CXCR3 siRNA-suppressed ECs (siCXCR3) to determine the role of CXCR3 in EC flow-mediated cellular responses. Treated cells are seeded into our flow chambers and either a pulsatile flow regimen applied (where shear starts and stops in a cyclical pattern and a small amount of backflow is noted; **Fig. 5**), or a steady-state laminar flow regimen applied (where shear was constant and unidirectional; **Supp. Fig. 4**) across time.

**Figure 5.**
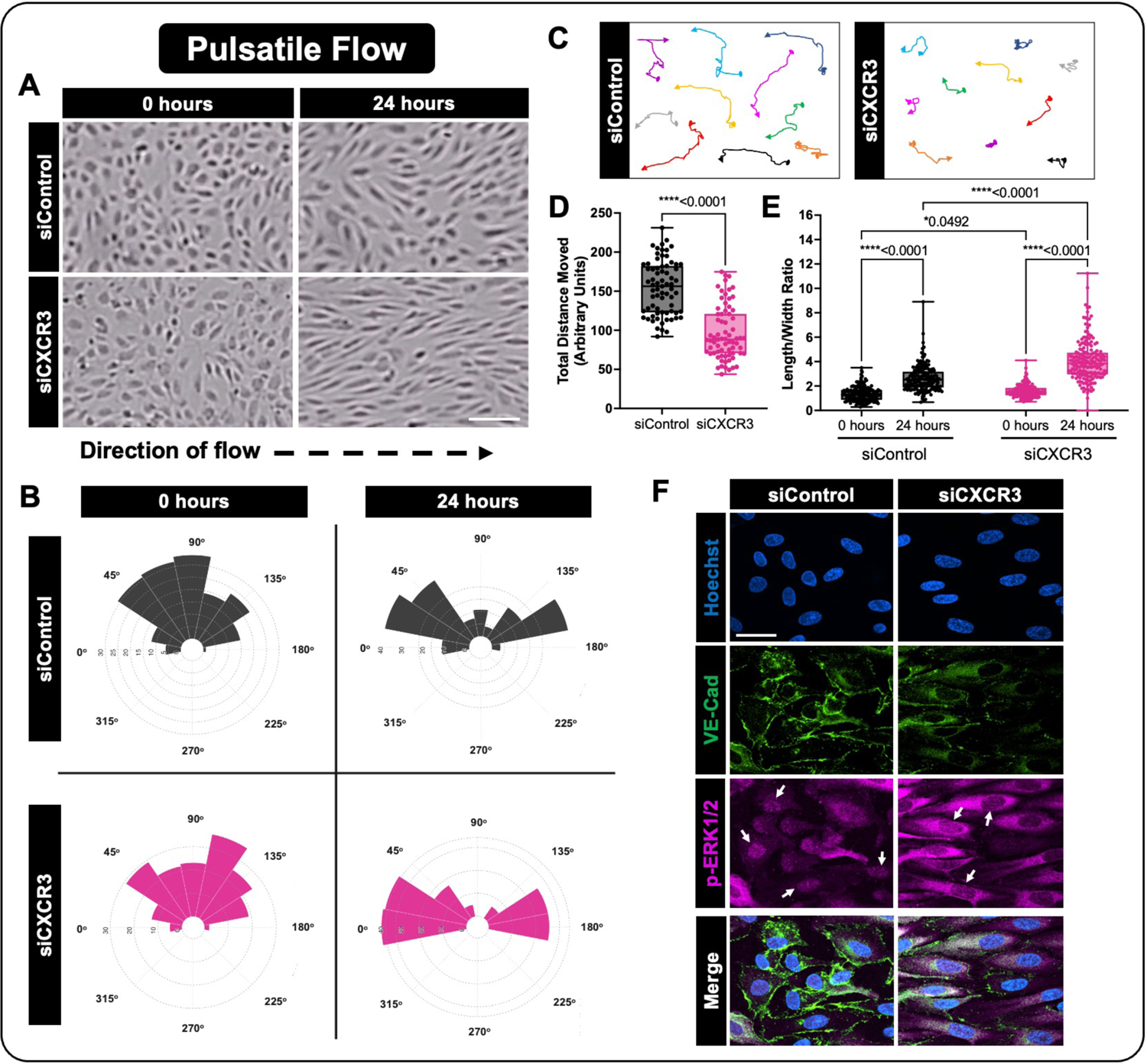
CXCR3 modulates EC motility and junction stability in response to a pulsatile flow regimen. **A)** Representative images of EC alignment to pulsatile flow at 0 and 24 hours, using ECs transfected with control (siControl) or siRNA against CXCR3 (siCXCR3). Scale bar represents 100µm. **B)** Distribution angles of the siControl (grey) or siCXCR3 (magenta) EC’s longest axis relative to the direction of flow at 0 and 24 hours. **C, D)** Analysis of siControl or siCXCR3 motility under pulsatile flow for 24 hours. Representative cell tracks and arrowheads represent the direction of cellular movement; circles represent the starting point of individual cells (C). Quantification of total distance of movement from the cells origin (D). **E)** Quantification of the length/width ratio of siControl or siCXCR3 ECs under pulsatile flow at 0 and 24 hours. **F)** Representative immunostaining images of siControl or siCXCR3 ECs treated with pulsatile flow for 24 hours. VE-Cadherin (green); p-ERK1/2 (magenta); nuclei (blue; Hoechst). White arrows indicate nuclear localization of p-ERK1/2 in ECs; Scale bar represents 50µm. For panels D and E, each dot represents an individual cell. Statistics for panel D were calculated using a Mann-Whitney test, and for panel E were calculated using a Kruskal-Wallis test with Dunn’s multiple comparisons test (omnibus p-value is <0.0001). Data are presented as box plots that display the median value with 1^st^ and 3^rd^ quartiles and min/max bars. All panels represent data from at least 3 independent experiments.

Under the pulsatile flow regimen, siControl ECs, which start as a randomly oriented cell population at time 0 (**Fig. 5A,B**), begin to move and reorganize, elongating their cell bodies, and ultimately arranging themselves to be obliquely oriented to the direction of flow by 24 hours (**Fig. 5A,B**; ∼45°). In the siCXCR3-treated ECs, the cells also begin as randomly oriented at time 0 (**Fig. 5A,B**), but become appreciably more elongated in the direction of flow (i.e. left to right, **Fig. 5A**), aligning nearly parallel to the pulsatile flow force by 24 hours (**Fig. 5A,B**; 0° and 180°). Interestingly, while under flow the siCXCR3 ECs move less distance from their origin than the siControl ECs (**Fig. 5C,D**) and become significantly more elongated over time (**Fig. 5E**). These same studies were carried out with a laminar flow regimen (**Supp. Fig. 4**). Under unidirectional, continuous laminar flow, siControl ECs align parallel to the direction of flow, and move in the direction of the flow force while elongating their cell bodies (**Supp. Fig. 4A-E**). In this case, siCXCR3 ECs directionally align identically to the siControl ECs (**Supp. Fig. 4A,B**), but still move less distance from their origin (**Supp. Fig. 4C,D**) and become significantly more elongated over time (**Supp. Fig. 4E**). Together, these data suggest that CXCR3 is used in ECs to interpret flow forces to induce motility and cellular alignment.

To understand how CXCR3 suppression could be affecting the endothelial response to flow, we immunostained cultures to assess proteins regulating EC motility (p-ERK1/2) and junctional stability (vascular endothelial cadherin; VE-Cadherin) ^31,32^. As mentioned, siCXCR3 cells reorient to be parallel to the direction of flow in response to pulsatile/interrupted flow, much more efficiently than their siControl counterpart cells, which can be easily seen by looking at nuclear orientation in these samples (**Fig. 5F**; Hoechst).

VE-Cadherin is decreased at EC-EC junctions in siCXCR3 ECs compared to siControl ECs (**Fig. 5F**; VE-Cad). Interestingly, while siCXCR3 ECs display higher levels of p-ERK1/2 immunolabeling overall (**Fig. 5F**; p-ERK1/2) the majority of the p-ERK1/2 signal is cytosolic, unlike the nuclear localization in siControl ECs (white arrows) (**Fig. 5F**). This is consistent with the p-ERK1/2 immunolabeling we see in the zebrafish cranial vasculature (**Fig. 4G,H**).

### CXCR3 is required *in vitro* and *in vivo* to promote pericyte recruitment

To functionally address how CXCR3-CXCL11 signaling can affect vasculature formation, we turned to type 1 collagen 3D tube formation assays where we can model vascular formation and pericyte recruitment (**Fig. 6A**) ^33-35^. In this assay, we seed ECs and pericytes in a 5:1 ratio in a type I collagen gel, with a serum free defined culture media. The ECs are allowed to form lumen-containing tubes and recruit pericytes over a period of 72 hours. Using this assay, we can manipulate either cell type to determine consequences on the opposing cell type over time ^33-35^. To begin, the 3D assays show that treatment with AMG487—the CXCR3 inhibitor—leads to expansion of EC tubes, while treatment with VUF11222—the CXCR3 activator—lead to inhibition of EC tube formation (**Supp. Fig. 5**). This was consistent with data noted *in vivo* (**Fig. 4**). Using neutralizing antibodies, we show that neutralization of CXCR3 or CXCL11 leads to an expansion in EC tube area independent of the presence of pericytes (**Fig. 6 B,C**), and a decrease in pericyte association with these EC tubes (**Fig. 6 D,E**; black arrows indicate pericytes that are not associated; white arrow heads indicate associated pericytes). To determine which cell type is driving this phenotypic response, we used siRNA to suppress CXCR3 (siCXCR3) or CXCL11 (siCXCL11) specifically in ECs. As shown, we can recapitulate the neutralizing antibody and pharmacologic phenotype of increased EC tube area by suppressing CXCR3 or CXCL11 in ECs only (**Fig. 6F**). Additionally, EC siCXCR3 or siCXCL11 treatment leads to EC tubes lacking pericyte association (**Fig. 6G**). Tracking the motility of the pericytes in these 3D assays, we show that in the absence of functional CXCR3 or CXCL11 activity, pericyte motility is suppressed, with the cells moving less distance from their origin (**Fig. 6H**).

**Figure 6.**
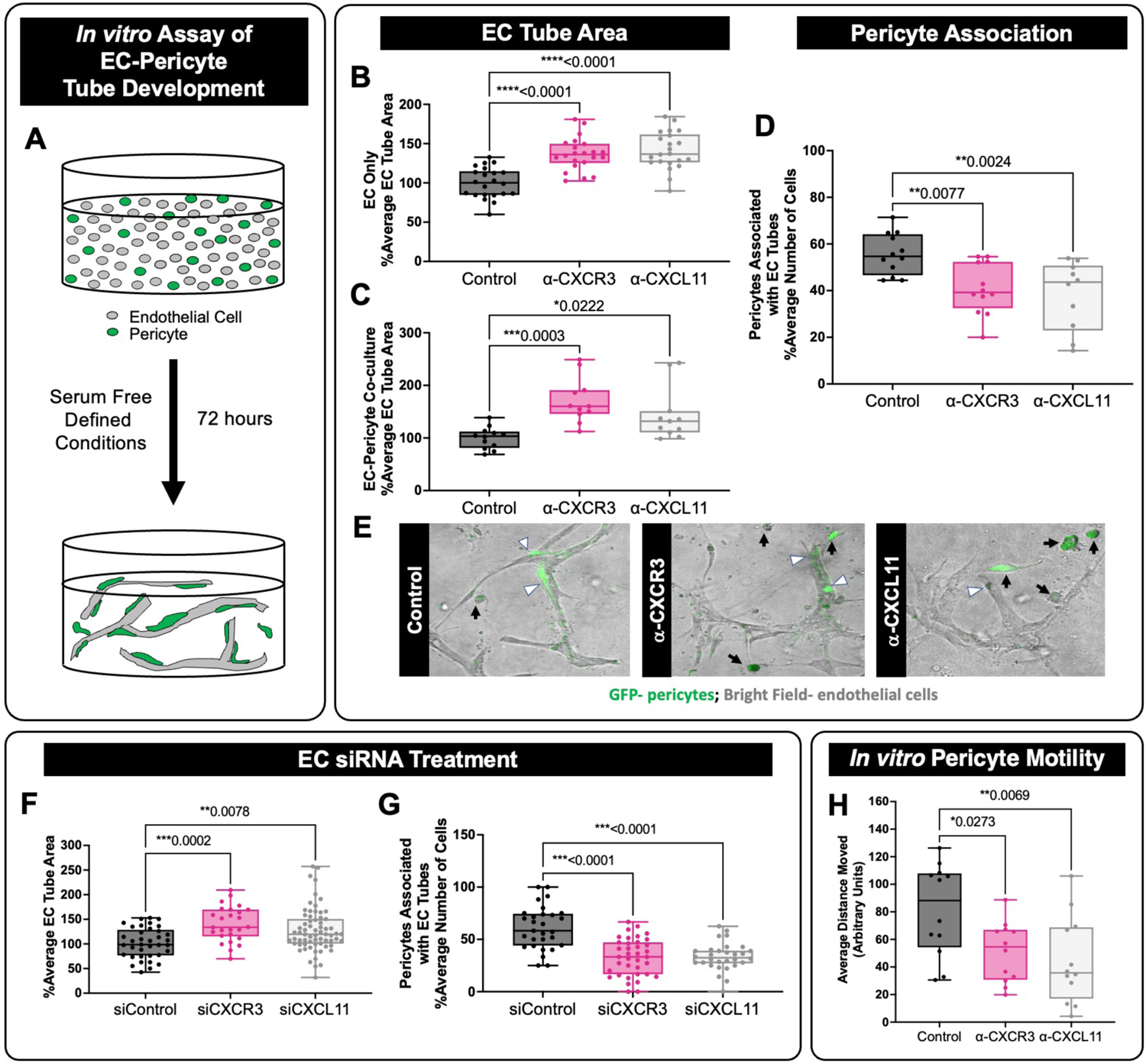
EC CXCR3 and CXCL11 suppress EC tube formation and pericyte recruitment *in vitro*. **A)** Schematic of 3D collagen type I EC-pericyte tube assembly assays. ECs and pericytes are used in a 5:1 ratio, and the pericytes are GFP-labeled for visualization. **B,C)** EC % average tube area measured from assays without (B) and with (C) pericytes present. Cultures were treated with neutralizing antibodies to CXCR3 (α-CXCR3) or to CXCL11 (α-CXCL11) and cultures were then allowed to assemble tubes for 72 hours. Neutralization of CXCR3 or CXCL11 leads to expanded EC tube area regardless of the presence or absence of pericytes. **D)** When pericytes are present, neutralization of CXCR3 or CXCL11 leads to reduced pericyte association with EC tubes. **E)** Representative image of the 3D EC-pericyte co-cultures at 72 hours. ECs are shown by bright field, and pericytes are GFP-labeled. White arrow heads indicate pericytes associated with EC tubes, black arrows indicate pericytes that are not associated with EC tubes. **F,G)** To determine if these effects are EC autonomous, we utilized siRNA to suppress CXCR3 (siCXCR3) or CXCL11 (siCXCL11) only in ECs, then incorporated these ECs into the 3D collagen gel assay with wild type pericytes. EC tube area is increased in siCXCR3 and siCXCL11 conditions compared to the siControl condition (F). Pericyte association with EC tubes is decreased following EC siCXCR3 or siCXCL11 treatment (G). All data are normalized to the control condition. **H**) Quantification of the distance that pericytes migrate in 3D collagen gel assays treated with neutralizing antibodies suggests that pericyte motility is impaired when CXCR3-CXCL11 signaling is inhibited. For panels B-D and F-H, each dot represents an individual 3D collagen assay. Statistics for panels B-D and H were calculated using one-way ANOVA with Dunnett’s multiple comparisons test. A Kruskal-Wallis test with Dunn’s multiple comparisons was performed for F and a Welch’s ANOVA test with Dunnett’s T3 multiple comparisons was performed for G. Omnibus ANOVA *P*-values (prior to the *post hoc* tests) are <0.0001 (B), 0.0006 (C), 0.0021 (D), 0.0003 (F), <0.0001 (G), and 0.0081 (H). Data are presented as box plots that display the median value with 1^st^ and 3^rd^ quartiles and min/max bars.

To assess pericyte recruitment *in vivo*, we used *Tg(fli1a:eGFP); Tg(pdgfrb: Gal4FF; UAS:RFP)* zebrafish embryos to mark the endothelium and *pdgfrb*-positive pericytes in the developing cranial vasculature, respectively. We treated the embryos with DMSO or AMG487 (to inhibit Cxcr3 activity) at 32 hpf and noted a suppression in the number of pericytes associated with the cranial vasculature at 72 hpf (white arrows, **Fig. 7A,B**). Using our TRAP-mRNA isolation methods (**Supp. Fig. 3**), we isolated EC-specific mRNA from DMSO control and AMG487 treated animals to assess EC-specific transcripts of target genes of interest. We were particularly interested in levels of *pdgfb* transcript, as this is a known regulator of pericyte recruitment and proliferation. As shown, transcript levels of *pdgfbb* and *pdgfab* are both suppressed in the endothelium of AMG487 treated zebrafish compared to DMSO treated control siblings (**Fig. 7C**), suggesting that loss of Cxcr3 activity leads to inhibition of pericyte recruitment via suppressed EC Pdgfb production. Together, these data provide *in vivo* evidence that is consistent with our *in vitro* results: CXCR3 is required for maximal *pdgfb* production in ECs and for pericyte recruitment to the vasculature.

**Figure 7.**
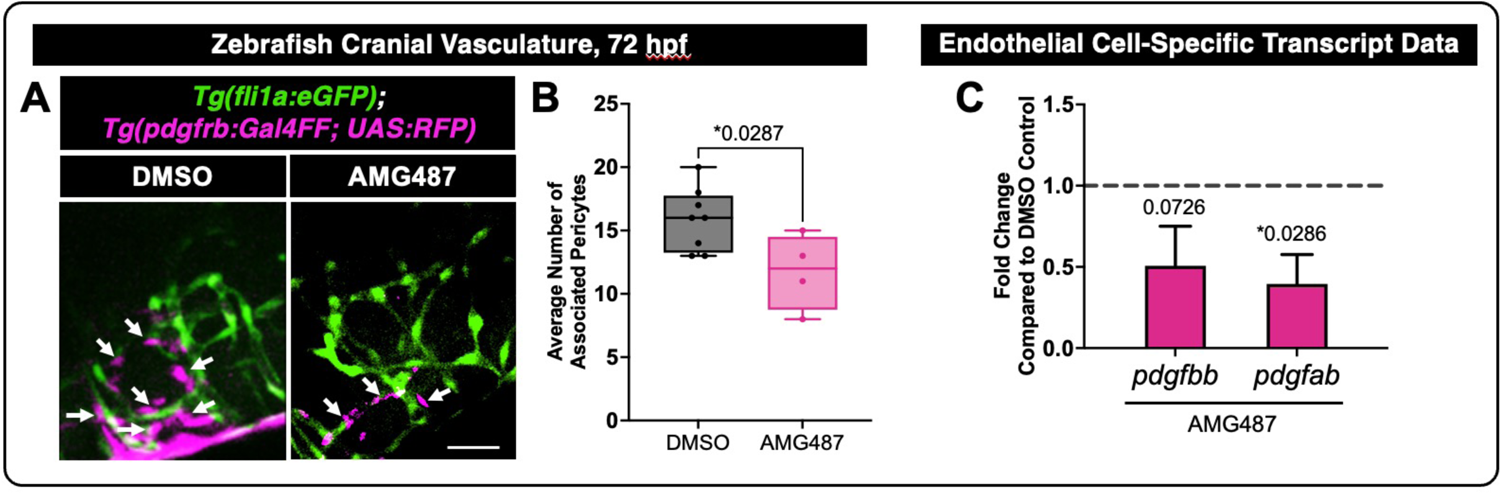
CXCR3 promotes EC transcription of *pdgfb* and for pericyte recruitment to blood vessels *in vivo*. **A)** Representative images from the *Tg(fli1a:GFP);Tg(pdgfrb:Gal4FF; UAS:RFP)* double transgenic zebrafish line marking endothelial cells in green and pericytes in magenta. Arrows point to pericytes that are associated with the cranial vasculature in 72 hpf zebrafish. DMSO or AMG487 (CXCR3 inhibition) was added at 32 hpf. Imaging is done in the zebrafish forebrain. Scale bar represents 50µm. **B)** Quantification of the number of pericytes associated with the vessels in the zebrafish forebrain following DMSO control or AMG487 treatment at 72 hpf. **C)** EC-specific mRNA was isolated using TRAP isolation protocols (Supp. Fig. 3) following DMSO control or AMG487 treatment. qPCR analysis for *pdgfbb* and *pdgfab* is shown as relative fold change compared to the DMSO control (grey dotted line). For panel B, each dot represents an individual zebrafish. Statistics for panel B were calculated using an unpaired t test, and one-sample t tests for panel C. Data in B are presented as box plots that display the median value with 1^st^ and 3^rd^ quartiles and min/max bars. Data in C are presented as bar graphs that display the mean±SD, from n=3 independent mRNA isolations.

## DISCUSSION

Here, we report a negative or delimiting role for CXCR3-CXCL11 signaling in the endothelium. We show that in the absence of CXCR3, ECs form poor junctions, have increased p-ERK1/2 levels, and, perplexingly, decreased cellular motility. In both the zebrafish cranial vasculature and our 3D modeling assays, ECs subsequently form tubes of a greater area, suggesting that physically larger ECs that have decreased motility and weaker junctions can allow for expansion of the vasculature. Additionally, following disrupted CXCR3-CXCL11 signaling, we see consistent loss of pericyte recruitment to EC tubes *in vitro* and *in vivo*, suggesting that regulation of this signaling pathway could underlie vessel stability at multiple stages of vascular development.

Historically, CXCR3 has been studied with a focus on its role in the immune response ^36-38^. It is highly expressed on effector T cells, and it is crucial for T cell trafficking and function. For example, CXCR3 is rapidly induced in naïve T cells following activation and remains preferentially expressed on Th1-type CD4+ T cells and effector CD8+ T cells ^39,40^. CXCR3 has also been shown to drive dendritic cell and a subset of B cell migration to the inflamed lymph node ^41,42^. Often overlooked, several studies have shown that CXCR3 is present in ECs of medium and large caliber vessels from different organs ^43^. In support, we find that early mesoderm precursors and endothelial cell populations express *cxcr3.1* in the zebrafish, suggesting a role for *cxcr3* isoforms in cardiovascular development.

Hemodynamic forces have a profound effect on cardiovascular development and play a crucial role in maintaining vascular integrity and function ^7,44,45^. Research into novel genes and molecular signaling pathways that can synchronize the mechanical and morphogenetic aspects of cardiovascular development are ongoing, and critical for fully understanding congenital vascular malformations ^46^. Here we show a biological role of CXCR3 in the context of vessel stabilization events during early cardiovascular development and demonstrate its functional role in interpreting flow forces to elicit changes in cellular behavior. In this study, we confirm CXCR3 as a blood flow-responsive gene and demonstrate that loss of CXCR3-mediated signaling impairs EC-EC junction formation and motility in response to pulsatile shear stress. These findings suggest that reductions in EC motility coupled with junctional impairment can lead to an expansion in vessel area. We also show that this expanded vasculature lacks normal pericyte association, which can lead to vascular instability over time.

Moving forward, CXCR3 signaling in ECs will be a novel pathway to consider when evaluating treatment strategies and therapeutic interventions aimed at controlling vascular disease and congenital vascular malformations. There is precedented for this concept, as other chemokine receptors, such as CXCR4, have been clearly demonstrated to be responsive to hemodynamic forces and can contribute to endothelial behavior. CXCR4 for instance, has been shown postnatally to be responsive to laminar shear stress, resulting in anti-atherogenic properties in ECs, altering survival, MCP-1, and IL-8 production ^47^. While these data may suggest secondary effects of the endothelium modulating the immune response, direct contributions of ECs or pericytes/smooth muscle cells are also likely contributing to altered vessel stability. In this work we have found a connection between EC-expressed CXCR3 and PDGFB production, leading to pericyte stability in the zebrafish cranial vasculature. Yet, how CXCR3 is connected to PDGFB transcription is entirely unknown. Moving forward, understanding the signaling nodes regulated by CXCR3 in ECs will be critical to elucidating its mechanistic role in vascular stabilization events.

So how is CXCR3 expression regulated in ECs during early cardiovascular development? Why is expression maintained at high levels when blood flow is suppressed? The KLF2 transcription factor is one of the best-studied blood flow regulated genes and plays an important role in regulating endothelial biology and vascular smooth muscle cell differentiation ^48,49^. Interestingly, KLF2 activity in immune cells has been shown to indirectly repress expression of pro-inflammatory chemokine receptors such as CXCR3 ^10^. For example, CD8 T cells that retain KLF2 expression fail to express CXCR3 and do not acquire the ability to migrate in response to CXCR3-CXCL10 activation ^50^. These data are compelling and suggests that KLF2 has the ability to suppress CXCR3 expression in cell-autonomous responses; however, to our knowledge, CXCR3 is not a direct KLF2-regulated target gene. In the vasculature, KLF2 expression decreases following impairment of blood flow, while we see an upregulation of CXCR3 in this context. Therefore, it is compelling to hypothesize then that KLF2 is a transcriptional repressor of CXCR3, though more work will need to be done in this area to explore this hypothesis.

For this work, we prioritized the use of pharmacologic tools to have the ability to assess both gain- and loss-of-function phenotypes present in the zebrafish following suppression or activation of CXCR3 activity at a discrete time point during development— coincident with the onset of blood flow. Moving forward, the development of genetic loss-of-function zebrafish models will be instrumental for understanding the role of CXCR3 in early mesenchyme lineage specification, vascular patterning events predating those we studied in this manuscript, and effects on other cardiovascular tissues.

## METHODS

### Zebrafish Methods and Transgenic Lines

Zebrafish *(Danio rerio)* embryos were raised and maintained as described ^51,52^. Zebrafish husbandry and research protocols were reviewed and approved by the Washington University Animal Care and Use Committee. All animal studies were carried out according to Washington University-approved protocols, in compliance with the *Guide for the Care and use of Laboratory Animals*.

Zebrafish transgenic lines—*Tg(fli1a:eGFP)^y1^; Tg(kdrl:mCherry-CAAX)^y171^; TgBAC(pdgfrb:Gal4FF)^ncv24^; Tg(5xUAS:RFP)^zf83^; Tg(kdrl:rpl10a-3xHA-2a-eGFP)^y530^*—are previously published ^21,53-56^.

### Reagents

AMG487 (CXCR3 inhibitor, #4487) and VUF11222 (CXCR3 activator, #5668) were purchased from Tocris Bioscience. Embryos were treated with 50µM of AMG487 or 25µM of VUF11222. BDM (2,3-Butanedione 2-monoxime, a myosin ATPase inhibitor) was purchased from Sigma-Aldrich (#B0753). Embryos were treated with 20 mM of BDM at 22 hpf and collected for translation ribosomal affinity purification (TRAP) at 48 hpf.

### Immunofluorescence staining

For immunofluorescence staining, cells were fixed in 4% paraformaldehyde for 30 min and permeabilized in ice-cold 100% methanol for 5 min. Cells were then blocked (5% BSA with 0.3% Triton X-100 in PBS) for 1h at room temperature. Primary antibody incubation was performed for 1h at room temperature in phospho-p44/42 MAPK (ERK1/2) (Thr202/Thr304) antibody (#4370, 1:200, Cell Signaling Technology) or human VE-Cadherin antibody (#AF938, 1:100, R&D Systems). Secondary antibody incubation was performed for 1h at room temperature at 1:2000 dilution using goat anti-Rabbit IgG (H+L) highly cross-adsorbed secondary antibody, Alexa Flour 594 (#A11037, Invitrogen) or donkey anti-Goat IgG (H+L) cross-adsorbed secondary antibody, Alexa Flour 488 (#A11055, Invitrogen). Nuclei were stained with Hoechst 34580 (#H21486, Invitrogen) for 15 min at room temperature. Imaging was performed using a W1 spinning disk confocal microscope (40x objective). Image analysis was performed in Fiji.

### 3D Collagen Matrix Assays and siRNA treatment

Human umbilical vein endothelial cells (HUVECs, Lonza) were cultured in growth media of M199 (#11150059, Gibco), 20% fetal bovine serum (FBS, Gibco), 0.05g heparin sodium salt (#H3393, Sigma-Aldrich), and 15 mg ECGS (#02-102, Sigma-Aldrich). Cells were routinely cultured on gelatin-coated plates at 37°C, 5% CO_2_. HUVECs were used from passages 2-6. Human brain vascular pericytes (HBVP, ScienCell) were cultured in 10% FBS in DMEM media on 1mg/ml gelatin coated flasks. HBVPs were used from passages 3-14. GFP-HBVPs were obtained from the G. Davis Lab (University of South Florida).

Small interfering RNA (siRNA) duplexes were synthesized by Life Technologies: CXCR3 siRNA (siCXCR3; #4392420, siRNA ID: s6014); CXCL11 siRNA (siCXCL11; #4392420, siRNA ID: s12620). Negative control siRNA (siControl; #4390847) was used as a control. The cells were transfected with 25 nM siControl or siCXCR3 using siPORT *NeoFX*™ Transfection Agent (#AM4511, Invitrogen) according to the manufacture’s protocol and twice across 3 days^33-35^.

Assays for 3-dimensional (3D) collagen type I gels were made using HUVECs treated with CXCR3 or CXCL11 siRNA co-cultured with HBVP-GFP. Culture media for the assays contained ascorbic acid (AA), FGF (#233-FB-025/CF, R&D systems), and IGF-2 (#292-G2-250, R&D systems). Additionally, FGF, SCF (#255-SC-010/CF, R&D systems), IL3 (#203-IL-010/CF, R&D systems), and SDF1a (#350-NS-010/CF) were supplied in the collagen gel ^33-35^. After 72 hours, the cells were fixed in PFA and images were taken with EVOS M5000.

### Imaging Analysis

Zebrafish imaging analysis and immunofluorescent imaging was done using a Nikon Ti2E microscope with a CSU-W1 spinning disk confocal (Yokogawa). Images were acquired using either a Nikon CFI plan Apo Lambda 20x objective or Nikon CFI plan Apo Lambda 60x objective. Zebrafish were anesthetized with MS-222 (#NC0872873, Western Chemical Inc.) and embedded in 0.8% low melting point agarose (#IB70056, IBI Scientific) for imaging. Live cell tracking was done using an EVOS M7000 microscope with a climate-controlled stage at 5% CO_2_ and 37°C.

### Flow Assays

HUVECs were transfected twice across 3 days with 25 nM of with control or CXCR3 siRNA. Cells were then seeded in µ-Slide I 0.4 Luer slide (#80176, ibidi USA Inc.) at 2.5 x 10^5^ cells/cm^2^ for 24 hours before the start of flow. Flow assays were carried out using an EVOS M7000 microscope (Invitrogen), equipped with an onstage incubator to maintain standard cell culture conditions. Time-lapse images were captured in bright field at 10x magnification (Olympus; UPlanSApo objective), every 20 minutes for 24 hours. The videos were created by stitching the 73 frames acquired together at a rate of five frames per second. A laminar flow regimen was applied at 15 dyn/cm^2^/s versus a constant pulsatile flow at 15 dyn/cm^2^/s and 30 RPM for 24 hours, using a modified Flocel pump and controller software (#WPX1). At the termination of the experiment, the cells were used for subsequent immunofluorescence staining applications ^30^.

### Translational ribosomal affinity purification (TRAP)

Zebrafish embryos were dechorionated with pronase (#10165921001, Roche). Embryos were deyolked with Deyolking buffer (55 mM NaCl, 1.9 mM KCl, 1.25 mM NaHCO_3_) containing protease inhibitor (#P8340, Sigma-Aldrich) and washed with a deyolking wash buffer (10 mM Tris-HCl [pH 8.5], 110 mM NaCl, 3.5 mM KCl, 2.7 mM CaCl_2_). Dechorionated and deyolked embryos were dounce homogenized in homogenization buffer (50 mM Tris-HCL [pH7.4], 100 mM KCl, 12 mM MgCl_2_, 1% NP-40) containing 1mM DTT (#646563, Sigma-Aldrich), 200 U/mL RNasin (#N2111, Promega), 100 µg/mL cycloheximide (#7698, Sigma-Aldrich), 1 mg/mL heparin (H3393, Sigma-Aldrich), and protease inhibitors. Homogenates were incubated with Dynabeads Protein G (#10007D, Invitrogen) coated with 3 µg of HA-Tag monoclonal antibody (#26183, ThermoFisher Scientific) at 4°C for overnight. Unbound materials were washed off using high salt homogenization buffer (50mM Tris pH 7.4, 300mM KCl, 12 mM MgCl_2_, 1% NP-40, 1 mM DTT, protease inhibitors, 200 U/mL RNAsin, 100 µg/mL cycloheximide, 1 mg/mL heparin). The mRNA was extracted from the Dynabeads and DNase treated using DNA/RNA/Protein extraction kit (#IB47702, IBI Scientific) and analyzed by RNA sequencing or qRT-PCR.

### RNA Extraction and qPCR Analysis

The embryos were lysed and purified with DNA/RNA/Protein extraction kit (#IB47702, IBI Scientific) and then cDNA generated with SuperScript™ IV VILO™ Master Mix (#11766050, Invitrogen) according to the manufacturer’s protocol. The TaqMan qPCR protocols were utilized to generate relative expression data, and analysis run using FAM channel of 96-well QuantStudio3 qPCR machine. For the quantification of mRNA levels, the ΔΔCT method was used. TaqMan probes: *cxcr3.1* (zebrafish, Dr03429972), *cxcr3.2* (zebrafish, APRWKAC), *cxcr3.3* (zebrafish, APPRRPE), *cxcl11.8* (zebrafish, APMF3GR), *pdgfbb* (zebrafish, APH6FCW), *pdgfab* (zebrafish, Dr03104374), *gapdh* (zebrafish, Dr03436842). All probes are FAM labeled (FAM; #4331182, Applied Biosystem). mRNA levels were normalized to *gapdh*.

### Statistics

Statistical analyses were performed using GraphPad Prism 10. Data normality was determined by using the D’Agostino-Pearson omnibus test. Statistical analyses, post-hoc tests and *P*-values are all described in corresponding figures and figure legends. Significance was determined by a *P*-value of 0.05 or less.

## Author Contributions

MG, JL, AL, ZB, GM, JA, SC, BC, YY, and ANS performed experiments; MG, JL, AL, ZB, YY, GM, BC, SC, and ANS analyzed results and made the figures; MG, JL, ZB, SC, and ANS designed the research and wrote the paper. Conflict-of-interest disclosure: The authors declare no competing financial interests. Correspondence: Amber N. Stratman, Department of Cell Biology and Physiology, Washington University School of Medicine St. Louis, MO, 63110;

## Acknowledgements

The authors would like to thank members of the Stratman laboratory for their critical comments on this manuscript. NIH/NIGMS R35 GM137976 (A.N.S.); Children’s Discovery Institute of Washington University and St. Louis Children’s Hospital (A.N.S.); the Washington University Institute of Clinical and Translational Sciences which is, in part, supported by the NIH/National Center for Advancing Translational Sciences (NCATS), CTSA grant #UL1TR002345 (A.N.S.).

## Supplemental Figure Legends

**Supplemental Figure 1.**
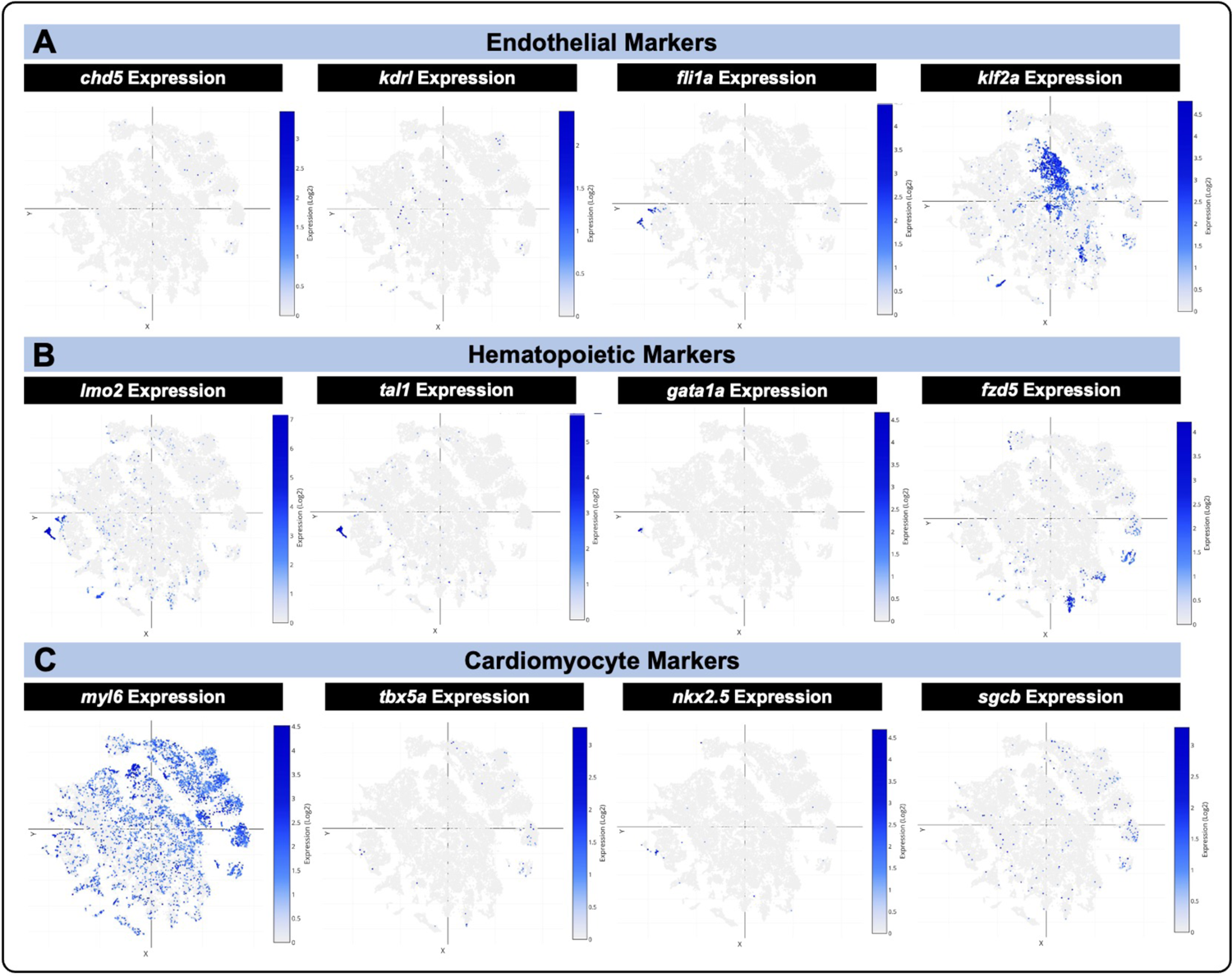
Single cell analysis of zebrafish embryos assessing localization of known endothelial, hematopoietic, and cardiomyocyte markers. UMAP plots of gene expression (blue dots) show where cells express known endothelial (A), hematopoietic (B), and cardiomyocyte (C) markers across all developmental stages present in this dataset ^25,26^. Darker blue dots equate to high gene expression, gray dots equate to no gene expression.

**Supplemental Figure 2.**
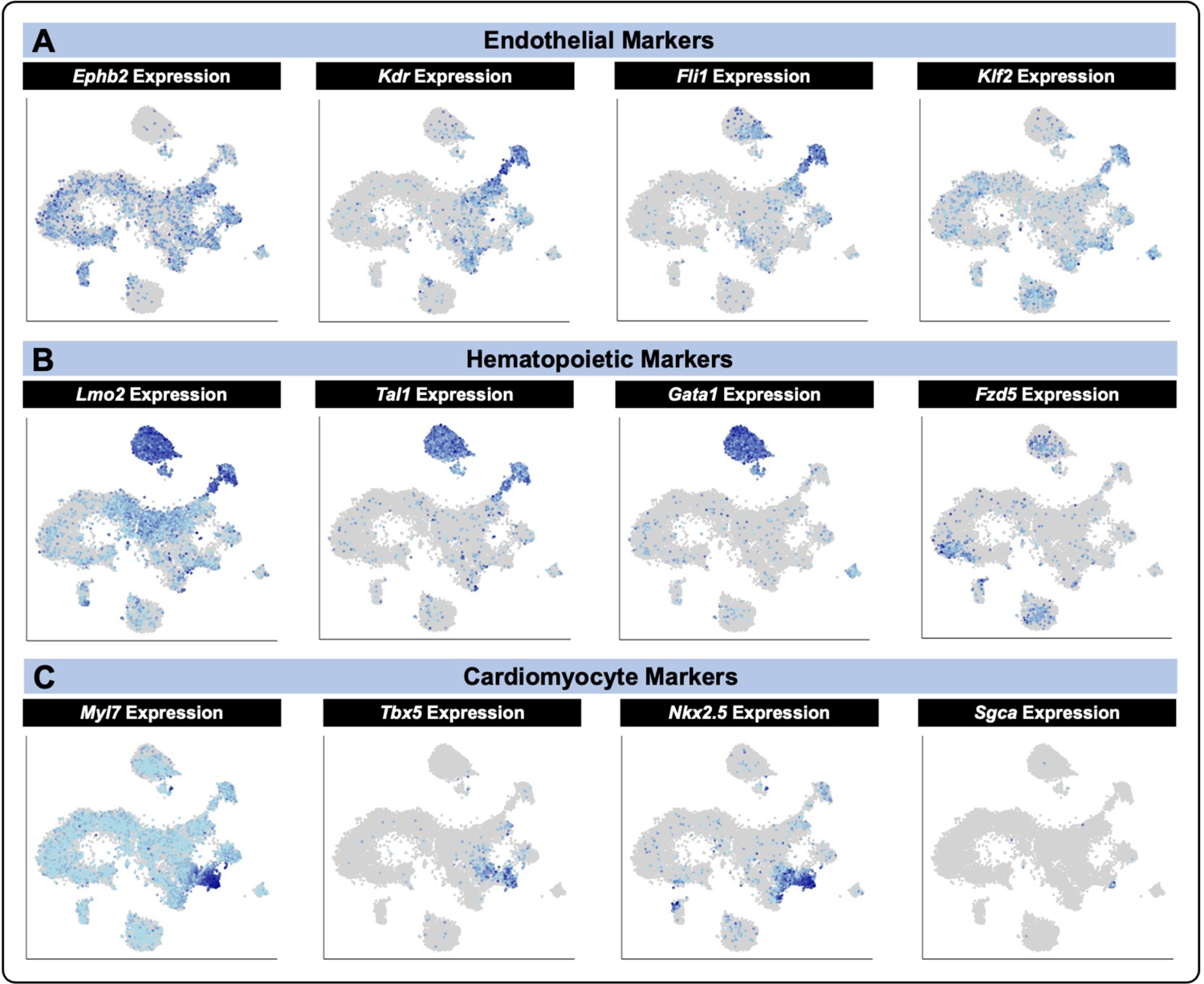
Single cell analysis of E8.25 mouse embryos assessing localization of known endothelial, hematopoietic, and cardiomyocyte markers. UMAP plots of gene expression (blue dots) show where cells express known endothelial (A), hematopoietic (B), and cardiomyocyte (C) markers at E8.25 in mouse development in this dataset ^27^. Darker blue dots equate to high gene expression, gray dots equate to no gene expression.

**Supplemental Figure 3.**
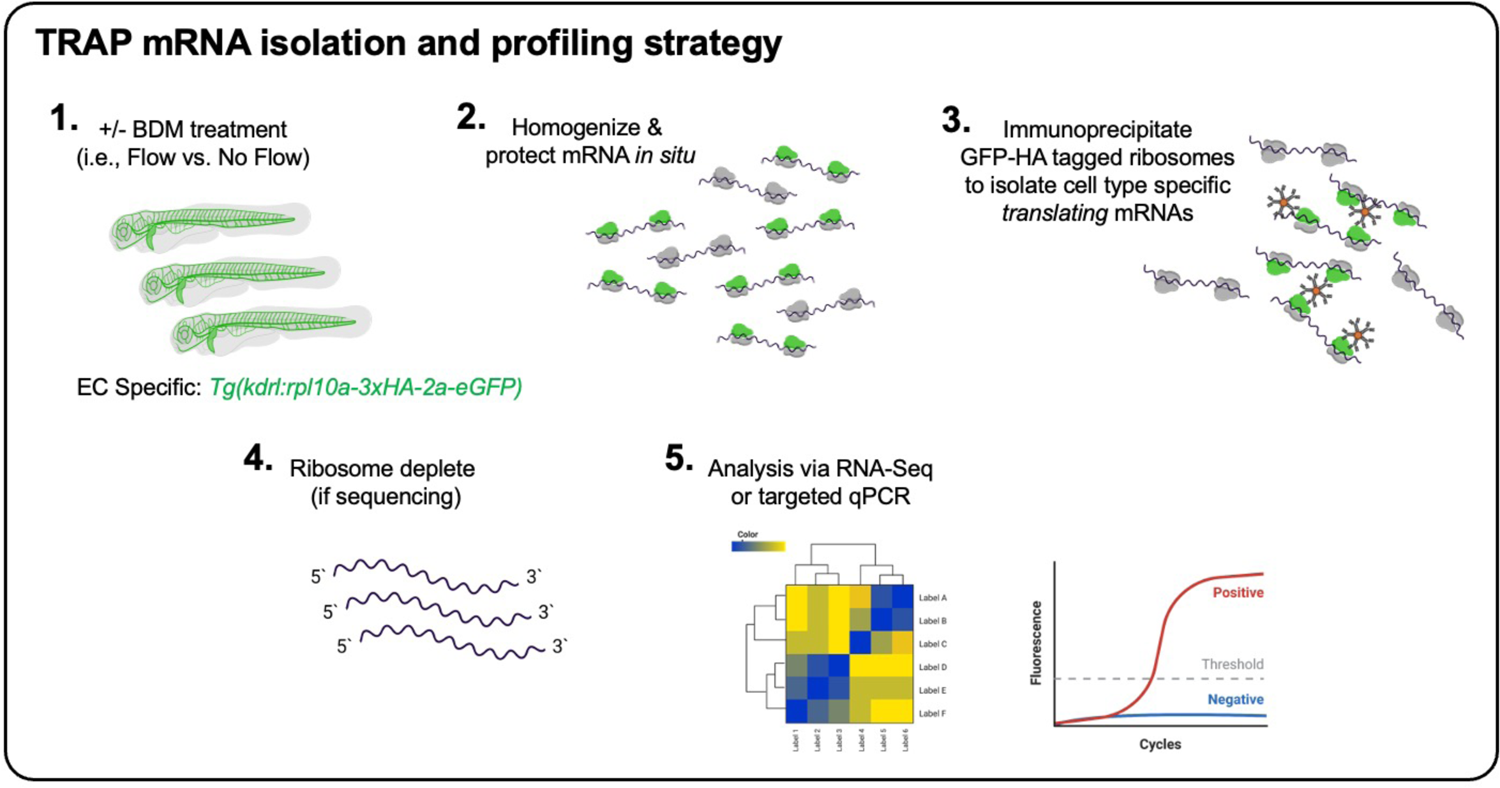
Schematic outline of the TRAP mRNA isolation and profiling strategy. The *Tg(kdrl:rpl10a-3xHA-2a-eGFP)* transgenic zebrafish line, which allows EC-specific tagging of ribosomes, was treated with 20 mM of BDM at 22 hpf to inhibit cardiac muscle contraction for flow/no flow mRNA profiling; or AMG487 for mRNA profiling following CXCR3 inhibition. Embryos were collected for immunoprecipitation of GFP-HA tagged ribosomes bound to translating mRNAs at 48 hpf. Ribosomal depletion from the samples is followed by analysis via mRNA-sequencing or qRT-PCR. This protocol allows for the isolation and profiling of EC-specific translating mRNAs that are collected from cells that remain in their native tissue environment.

**Supplemental Figure 4.**
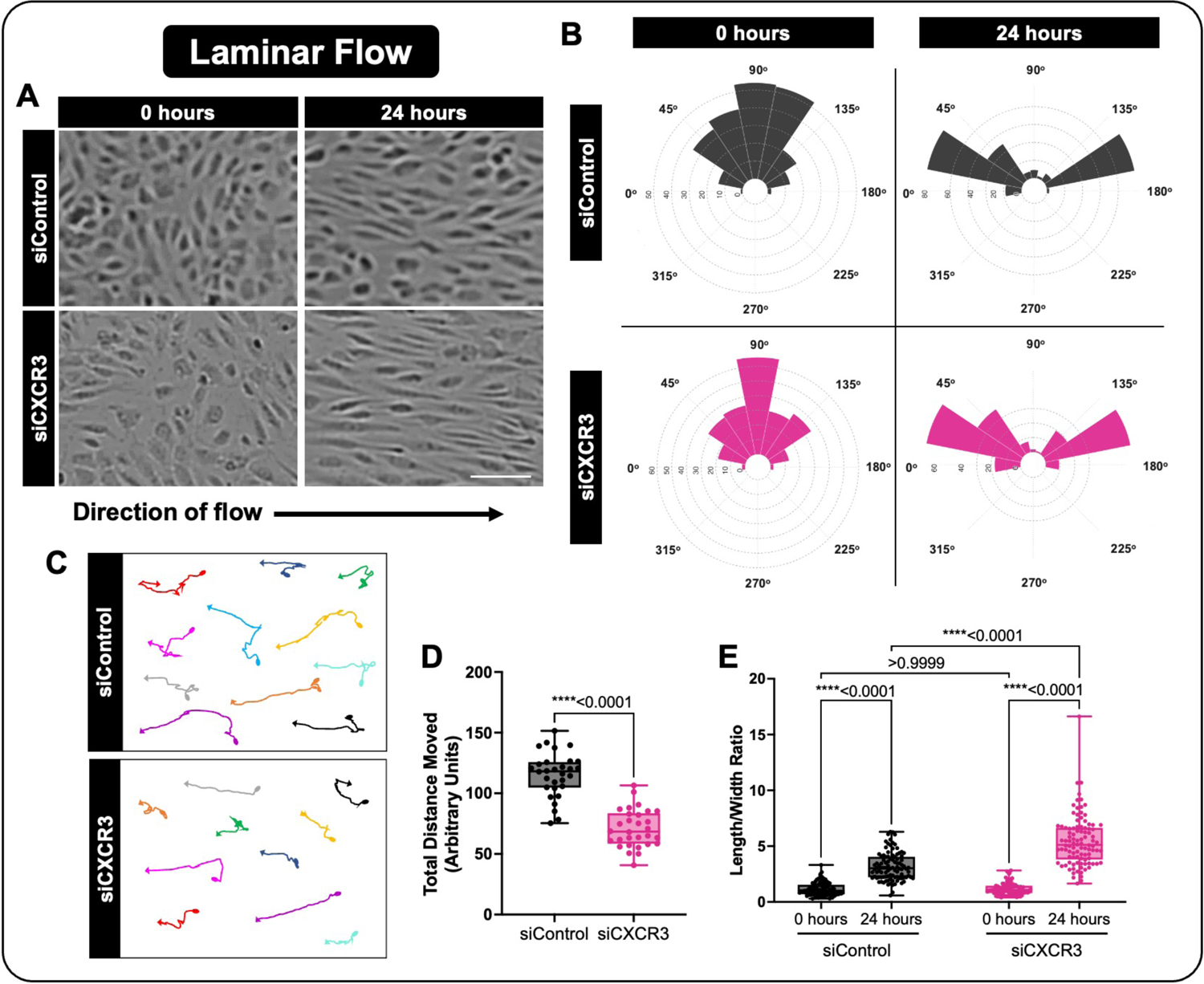
CXCR3 modulates EC motility and elongation in response to a laminar flow regimen. **A)** Representative images of EC alignment to laminar flow at 0 and 24 hours, using ECs transfected with control (siControl) or siRNA against CXCR3 (siCXCR3). Scale bar represents 100µm. **B)** Distribution angles of the siControl (grey) or siCXCR3 (magenta) EC’s longest axis relative to the direction of flow at 0 and 24 hours. **C, D)** Analysis of siControl or siCXCR3 motility under laminar flow for 24 hours. Representative cell tracks and arrow heads represent the direction of cellular movement; circles represent the starting point of individual cells (C). Quantification of total distance of movement from the cells origin (D). **E)** Quantification of the length/width ratio of siControl or siCXCR3 ECs under laminar flow at 0 and 24 hours. For panels D and E, each dot represents an individual cell. Statistics for panel D were calculated using an unpaired t test, and for panel E were calculated using a Kruskal-Wallis test with Dunn’s multiple comparisons test (omnibus p-value is <0.0001). Data are presented as box plots that display the median value with 1^st^ and 3^rd^ quartiles and min/max bars. All panels represent data from at least 3 independent experiments.

**Supplemental Figure 5.**
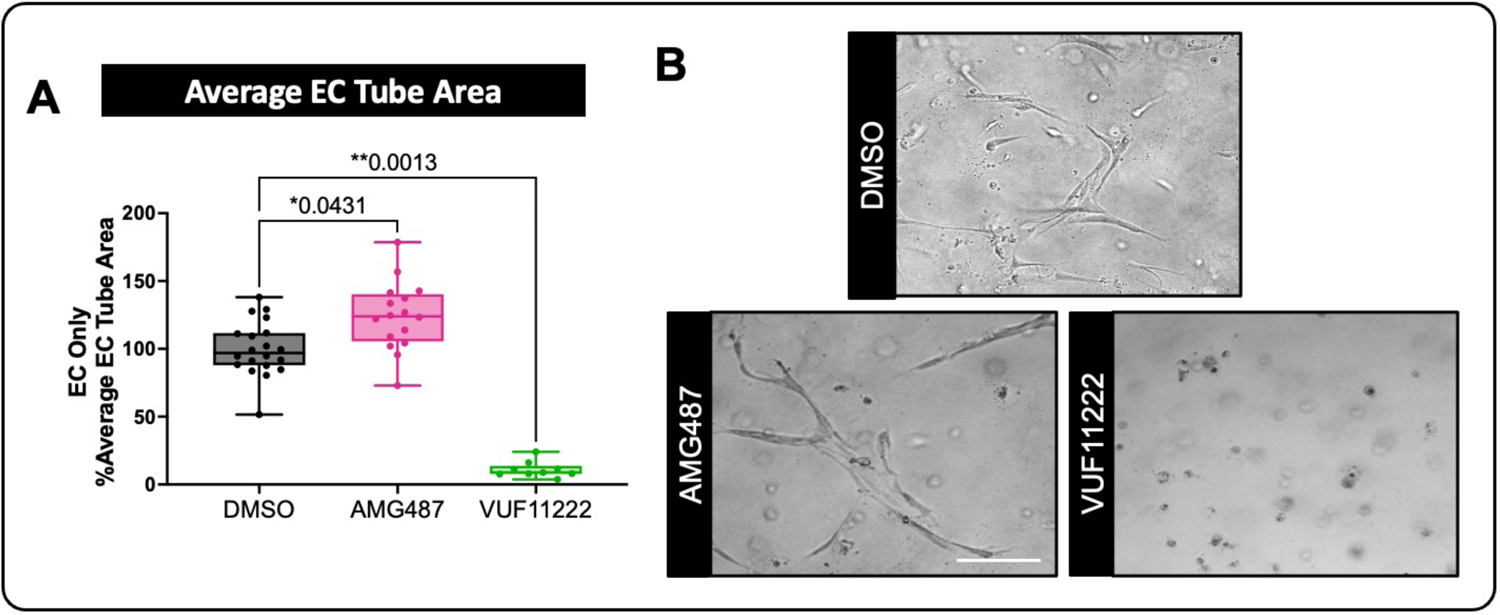
Modulation of CXCR3 activity by pharmacologic reagents alters EC tube formation *in vitro*. **A)** EC % average tube area in 3D collagen I matrix assays (with ECs only) following treatment with a pharmacological CXCR3 inhibitor (AMG487) or activator (VUF11222) versus DMSO as a control. **B)** Representative images of EC tubes within the 3D collagen I matrix assays *in vitro*. Scale bar represents 100µm. For panel A, each dot represents an individual 3D collagen assay. Statistics for panel A were calculated using a Kruskal-Wallis test with Dunn’s multiple comparisons (omnibus p-value is <0.0001). Data are presented as box plots that display the median value with 1^st^ and 3^rd^ quartiles and min/max bars.

